# Somatotopic specificity of perceptual and neurophysiological changes associated with visuo-proprioceptive realignment

**DOI:** 10.1101/2020.07.10.197632

**Authors:** Jasmine L. Mirdamadi, Courtney R. Seigel, Stephen D. Husch, Hannah J. Block

**Affiliations:** Program in Neuroscience, Indiana University Bloomington; Department of Kinesiology, School of Public Health, Indiana University Bloomington

## Abstract

Spatial realignment of visual and proprioceptive estimates of hand position can occur in response to a perturbation. E.g., viewing the hand underwater while washing dishes. This form of multisensory perceptual learning presumably affects perceived hand position, which is used in movement planning. Consistent with this idea, we recently observed changes in motor cortex (M1) index finger representation associated with visuo-proprioceptive misalignment at that fingertip (Munoz-Rubke et al., 2017). In three experiments with healthy human participants, we asked whether these changes are specific to the brain’s representation of the misaligned finger (somatotopically focal), or whether they extend to other parts of the hand and arm that would be needed to move the misaligned finger (somatotopically broad). In Experiments 1 and 2, participants experienced misaligned or veridical visuo-proprioceptive information about the index fingertip. Before and after the perceptual alignment task, we used transcranial magnetic stimulation (TMS) to assess M1 representation of five hand and arm muscles. The index finger representation showed the expected association between M1 excitability and visuo-proprioceptive realignment, as did the pinkie finger representation to a lesser extent. Forearm flexors, forearm extensors, and biceps did not show any such relationship. In Experiment 3, we asked subjects to indicate their proprioceptive estimate of the fingertip, knuckle, wrist, and elbow, before and after misalignment at the fingertip. Proprioceptive realignment at the knuckle, but not the wrist or elbow, was correlated with realignment at the fingertip. These results suggest the effects of perceptual perturbation are somatotopically focal in both sensory and motor domains.

**Significance Statement:** Multisensory and motor processing have largely been examined separately, limiting our understanding of how these systems interact. Motor adaptation is thought to affect somatosensory perception and neurophysiology, but the reverse interaction, an effect of perceptual learning on the motor system, has rarely been considered. Here we examine the effect of a somatotopically localized visuo-proprioceptive perturbation on motor neurophysiology and conscious perception. We found somatotopically-focal effects in both domains. This correspondence highlights the tight relationship between sensory and motor systems, but also raises the possibility that perceptual learning may not generalize to other body parts as motor learning does. Rather, it appears to create a localized distortion in the multisensory body representation, with correspondingly local changes in the motor cortex representation.

## Introduction

The brain normally has access to true hand position (*Y*) through vision and proprioception. The image of the hand on the retina provides a visual estimate 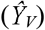, while receptors in the muscles and joints of the arm and hand provide a proprioceptive estimate 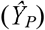. To form a single estimate with which to guide behavior, the brain is thought to weight and combine them into a single, integrated estimate of hand position (Ghahramani et al. 1997). If there is a spatial misalignment between modalities, as when the hand is submerged in water, which refracts light (Fig. 1A), the brain can realign one or both 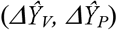 to compensate (Block and Bastian 2011; Ghahramani et al. 1997). Such a change in multisensory perception presumably affects perceived hand position, and thus motor planning with that hand. However, multisensory and motor processing have largely been examined separately, limiting our understanding of how these systems interact.

**Figure 1.**
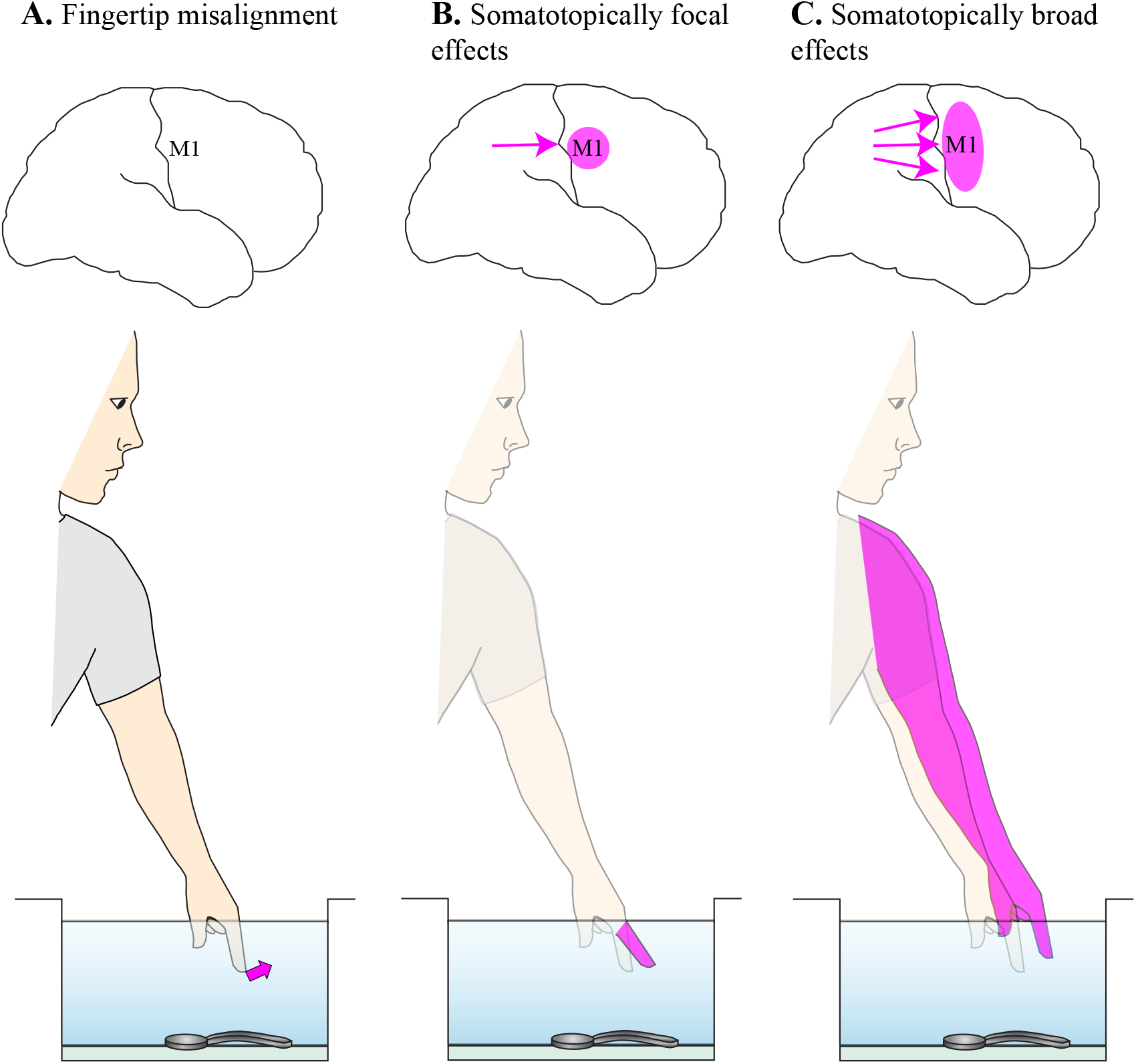
Visuo-proprioceptive representation of the body parts is both critical for motor control and highly plastic. **A.** Viewing the finger underwater shifts visual information about the finger away from proprioceptive information (pink arrow). The brain realigns visual and proprioceptive estimates of finger position, resulting in an integrated estimate closer to the visual distortion. The new estimate is presumably used in motor planning, and the realignment process is associated with excitability changes in the M1 representation of that finger (Munoz-Rubke et al. 2017). However, the somatotopic specificity is unknown. **B.** One possibility is that such changes are somatotopically focal, affecting primarily the misaligned body part (pink) and thus distorting the body schema nearby. Parietal changes associated with realignment are expected to affect only the finger area of M1 in this case (arrow). **C.** An alternative possibility is that the brain generalizes such changes more broadly, perhaps including other M1 representations needed to move the misaligned effector. In this case, parietal changes would be expected to affect more parts of M1 (arrows).

Motor learning is known to affect sensory perception (Ostry and Gribble 2016; Sexton et al. 2019), but the reverse interaction, an effect of perceptual learning on the motor system, has rarely been considered. If perceptual learning affects motor neurophysiology, we might expect to see a change in primary motor cortex (M1). M1 is closely interconnected with somatosensory cortex (SI), and somatosensory responses have been recorded in M1 neurons (Hatsopoulos and Suminski 2011). Proprioceptive training or stimulation has been found to affect motor learning (Wong et al. 2012) or networks (Carel et al. 2000; Lewis and Byblow 2004; Vahdat et al. 2014), respectively. However, such studies have largely been limited to the somatosensory system, with multisensory contributions to hand perception often overlooked.

We recently assessed M1 excitability for an index finger representation with transcranial magnetic stimulation (TMS), before and after subjects experienced misaligned or veridical visuo-proprioceptive information about that finger. The results suggest M1 changes are modality-specific, with a decrease in M1 excitability associated with proprioceptive realignment, and an increase in M1 excitability associated with visual realignment (Munoz-Rubke et al. 2017). Such modality-specific changes would suggest M1 is affected by plasticity in areas traditionally considered unisensory, such as SI (Krubitzer and Kaas 1990; Ostry and Gribble 2016) or early visual areas that have indirect connectivity with M1 (Strigaro et al. 2015).

The somatotopic specificity of changes in M1 due to perceptual learning is unknown, but may have functional significance. In other words, if misaligned visuo-proprioceptive information about the index finger is presented, would changes to M1 and to body perception be limited to the representation of an index finger muscle (somatotopically focal: Fig. 1B), or would representations of the entire effector (including biceps and forearm muscles) be affected (somatotopically broad: Fig. 1C)? A somatotopically focal interaction would be consistent with the effects of peripheral somatosensory stimulation on M1. For example, vibration of one muscle increases M1 excitability for that muscle, but decreases it for neighboring muscles (Rosenkranz and Rothwell 2004). However, a somatotopically broad interaction would be consistent with what we know of the generalization of motor learning within a limb. For example, when subjects learned a force production motor skill with their hand, it transferred to their arm and vice versa (Rajan et al. 2019).

Here we examined effects of a somatotopically localized visuo-proprioceptive perturbation on M1 neurophysiology and conscious perception. In both domains, we asked whether such effects are specific to the brain’s representation of the misaligned finger (somatotopically focal), or whether they extend to other parts of the hand and arm that would be needed to move the misaligned finger (somatotopically broad). In Experiments 1 and 2, we used TMS to measure M1 excitability for five hand and arm muscles, before and after subjects experienced misaligned or veridical visuo-proprioceptive information about the index fingertip. In Experiment 3, we asked subjects to indicate their proprioceptive estimate of four hand and arm joints, before and after misalignment at the fingertip, to assess the somatotopic focality of changes in perception.

## Methods

### Participants

Participants in all three experiments were healthy right-handed young adults, with similar age ranges and gender balances. 27 subjects (21.7 ± 3.3 years, mean ± SD, 16 males) participated in Experiment 1. 26 subjects (21.9 ± 4.7 years, 9 males) participated in Experiment 2. Fourteen subjects participated in Experiment 3 (21.8 ± 2.49 years, 8 males). All participants reported themselves to be neurologically healthy with normal or corrected to normal vision. All participants gave written informed consent. All procedures were approved by the Indiana University institutional review board.

### Experiments 1 and 2: M1 hand and arm representation

#### Sessions

Subjects completed two sessions each: the misaligned session (experimental) and the veridical session (control). In the misaligned session, a 70 mm misalignment between visual and proprioceptive information about the left index fingertip was gradually imposed. In the veridical session, visuo-proprioceptive information about the left index fingertip remained veridical throughout. The two sessions were separated by at least 5 days (mean 14.2 days, range 5-69 days) and session order was counterbalanced across participants.

In each session, TMS measurements were made pre- and post-perceptual alignment task (Fig. 2A). Experiments 1 and 2 were identical except for which M1 representations were targeted for TMS, and which muscles were recorded. In Experiment 1 (N = 27), TMS-evoked activity was recorded from two muscles in the left hand: first dorsal interosseus (FDI) and abductor pollicis brevis (ADM). In Experiment 2 (N = 26), TMS-evoked activity was recorded from three muscles in the left arm: flexor carpi radialis (FCR), extensor carpi radialis (ECR), and biceps brachii (BB). At the end of each session in both experiments, subjects were asked to rate their level of attention, fatigue, pain from TMS, and quality of sleep the night before, on a scale of 1-10, with 10 being the most attention, fatigue, or pain, and the best sleep.

**Figure 2.**
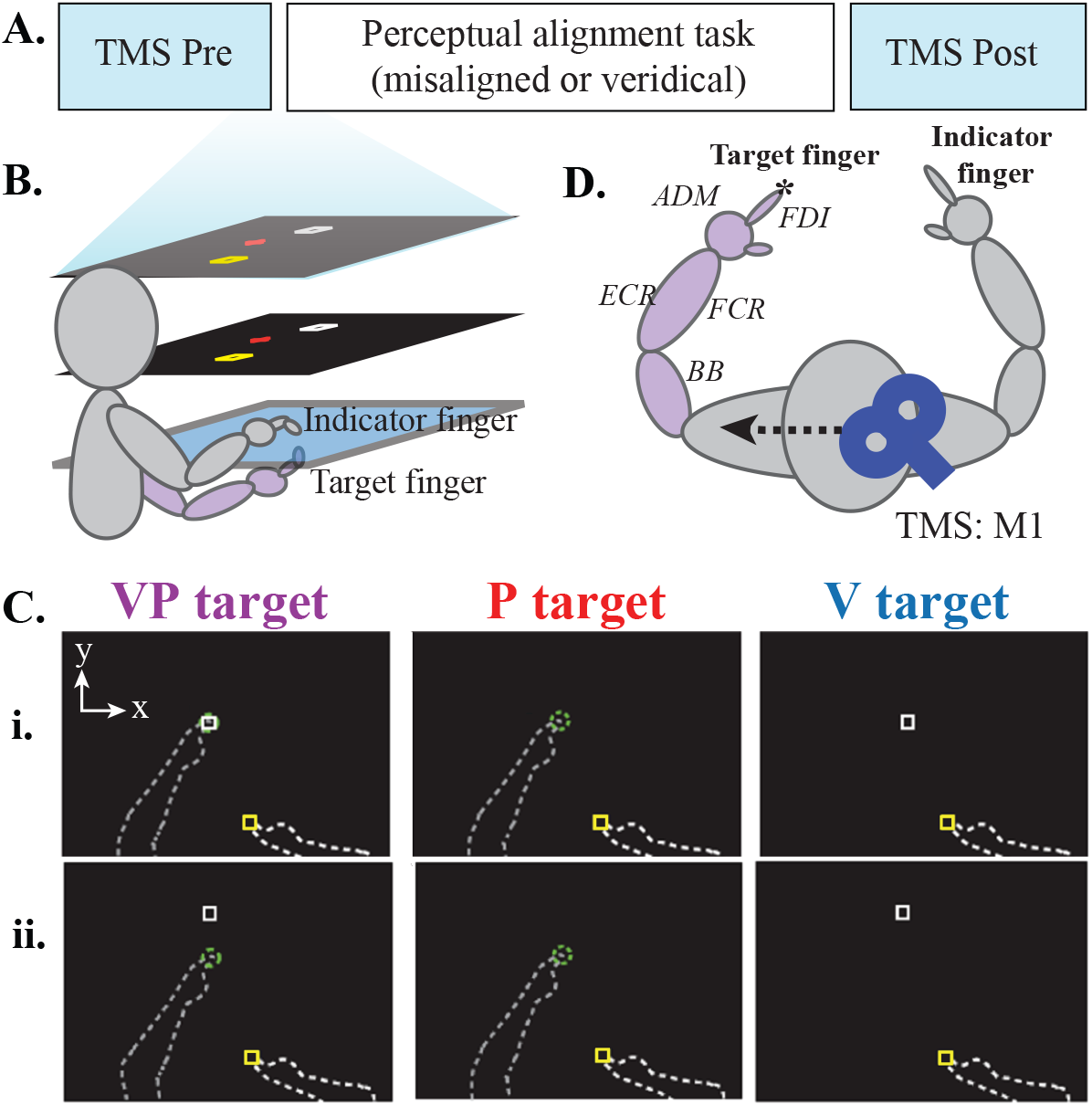
Experiment 1 and 2 methods. **A.** Session design. Each subject completed two sessions, distinguished by the VP target being veridical throughout (control) or gradually misaligned. **B.** Perceptual alignment task apparatus. Images viewed in the mirror (middle layer) appeared to be in the plane of the touchscreen glass (bottom layer). Subjects pointed with their right (indicator) index finger above the glass to where they perceived targets related to their left (target) index finger, which was positioned on a tactile marker beneath the glass (purple). **C.** Bird’s-eye view of the three target types. Indicator finger began in the yellow start box and moved to where the subject perceived a white box (V target), their left fingertip (P target), or both together (VP target). Dashed lines not visible to subject. No performance feedback or knowledge of results, and subjects were instructed to take their time and be as accurate as possible. **i.** Veridical targets. The white square was projected directly over the target fingertip. **ii.** In the misaligned session only, the white box was gradually shifted forward from the target fingertip. **D.** TMS setup. TMS over the right M1 was used to assess I/O curves in the left, target limb (purple). This was done pre- and post-alignment task for the target hand FDI and ADM (Experiment 1) or the target arm FCR, ECR, and biceps (Experiment 2). *Misaligned visual information was presented at the target index fingertip.

#### Perceptual alignment task apparatus and setup

Participants were seated at a custom 2-D virtual reality apparatus (Fig. 2B) without vision of their arms. The apparatus contained a two-sided touchscreen consisting of two infrared touch overlays (PQLabs) with a 3-mm-thick pane of glass sandwiched between. This was positioned in the horizontal plane beneath the mirror, where subjects viewed visual information that appeared to be in the plane of the touchscreen.

We used a bimanual task in which subjects indicated with their right index fingertip (indicator finger) the position of three types of target (Fig. 2C) associated with the left index fingertip (target finger). The indicator finger remained above, and the target finger below, the touchscreen surface throughout the session. Visuo-proprioceptive (VP) targets consisted of the target fingertip, touching a tactile marker on the lower surface of the touchscreen, with a 1-cm white square projected on top. Visual-only (V) targets were indicated by the white square alone, with the target hand resting in the subject’s lap. Proprioceptive-only (P) targets consisted of the target index finger placed on a tactile marker on the lower surface of the touchscreen, with no white square. For P and VP trials, participants actively placed their target finger on the tactile marker. This active component is likely to increase the ratio of proprioceptive to visual realignment (Welch et al. 1979; Welch and Warren 1980) and encourage visuo-proprioceptive integration (Balslev et al. 2006). Participants were told that on VP trials, the white box would appear directly over their target fingertip and that they should aim to place their indicator finger at that location. To best measure perceptual changes about the target finger and prevent motor adaptation of the indicator hand, participants were instructed to reach at a comfortable pace and not to rush. No online or endpoint feedback about the indicator hand was given at any point in the task, and subjects had no direct vision of either hand. Thus, participants did not know anything about their accuracy in indicating the positions of the targets.

#### Perceptual alignment task procedures

At the start of every trial, an 8-mm blue cursor representing indicator finger position appeared to help subjects place their indicator finger in the correct start position, which was represented as a yellow square. Once the indicator finger entered the yellow start square, the blue cursor disappeared so that no feedback was given regarding the movement of the indicator finger during the trial. As in previous studies, there were two possible target position and five possible start locations (Block et al. 2013; Liu et al. 2018; Munoz-Rubke et al. 2017). This was intended to add variability to the movement direction and extent of the indicator finger. The goal was to encourage subjects to accurately indicate the perceived target position with their indicator finger, rather than make any stereotyped movements. A second goal was to make it less likely that subjects would make systematically larger reaches in the misaligned session compared to the veridical session, due to the visuo-proprioceptive mismatch. Throughout each trial, participants were told to focus on a red fixation cross. This was done to better capture changes in visual estimates of target position, as opposed to a change in eye position. The red cross appeared within a 10-cm zone around the target, with location randomized across trials. Reaching performance has been shown to be somewhat affected by gaze direction (Henriques et al. 2003), but sometimes subjects actually misunderstand the directions and attempt to place their indicator finger at the red cross, instead of at the target, especially on P targets where there is no other visual information. To evaluate whether this occurred, we computed a regression between red cross location and P target estimates (indicator finger endpoint position) for each session. For Experiment 1, this R^2^ was 0.053 (mean) ± 0.011 (95% CI) in the misaligned session and 0.050 ± 0.014 in the veridical session. For Experiment 2, this R^2^ was 0.050 ± 0.014 and 0.024 ± 0.006 in the misaligned and veridical sessions, respectively. These low values suggest that subjects’ target estimates were not likely influenced by the red cross position.

The perceptual alignment task portion of each session began with a 69-trial baseline V, P, and VP trials, with the white square presented veridically over the target fingertip on VP trials (Fig. 2Ci). The main portion consisted of 84 trials (42 VP, 21 V, and 21 P) presented in alternating order (VP, P, VP, V, etc). In the veridical session, target presentation was also veridical. However, in this portion of the misaligned session, the position of the white square was gradually shifted forward from the target finger in the positive y direction (away from the subject’s body), 1.67 mm per VP trial (Fig. 2Cii). Thus, after 84 trials, there was a total misalignment of 70 mm between visual and proprioceptive information about the target fingertip. Most participants did not notice this gradual perturbation. At the end of each session, participants were asked “Did it feel like the white box was always on top of your target finger?” If they responded that it did not, we asked “In a particular direction, or all over the place?” If they indicated a particular direction, we asked them to estimate the magnitude. In Experiment 1, three participants reported a perceived forward displacement in the misaligned session only, one in the veridical session only, and one in both sessions. In Experiment 2, four participants reported a perceived forward displacement in the misaligned session only, and three reported this in the veridical session only. Because this is consistent with our previous use of this paradigm, we did not exclude any of these subjects (Block et al. 2013; Munoz-Rubke et al. 2017).

#### Perceptual alignment task analysis

Spatial realignment of visual and proprioceptive estimates of target finger position were taken from V and P trials, respectively. Proprioceptive realignment is expected to manifest as overshooting of the P target by the end of the misaligned VP trials, relative to the beginning; i.e., the subject comes to feel his target finger is further away than it is as the gradual misalignment of VP targets occurs. Visual realignment is expected to manifest as undershooting of the V target by the end of the misalignment phase, relative to the beginning. I.e., the subject comes to feel the visual representation is closer than it really is. As we have done previously (Block et al. 2013; Block and Bastian 2011, 2012), we quantified visual and proprioceptive realignment 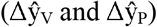 by subtracting indicator finger endpoint positions on the first four V or P trials of the misalignment phase from the last four:

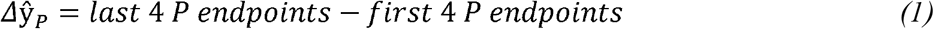

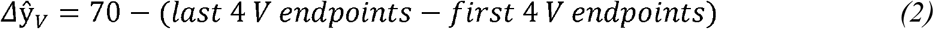

Realignment was computed relative to actual target position, which does not change for P targets but shifts forward 70 mm for V targets during the misaligned session. Thus, for both modalities, realignment in the expected direction comes out positive. These values were computed for the veridical session as well, using the first and last 4 V and P trials of the main 84 trials just like the misaligned session.

#### TMS setup and procedures

Before and after the perceptual alignment task in each session, TMS was used to measure the right hemisphere motor cortex excitability (Fig. 2D); i.e., the hemisphere controlling the left (target) hand that experiences visuo-proprioceptive misalignment in the misaligned session but performs no motor task other than touching the tactile marker. During TMS, both arms rested on a pillow in the subject’s lap. TMS was delivered using a Magstim 200^2^ stimulator (Magstim Company LTD, UK) with a 70 mm figure-of-eight coil. The coil was positioned tangential to the skull with the handle pointing posteriorly, 45□ to the inter-hemispheric line to evoke posterior-to-anterior current in the cortex (Rossini et al., 2015). We used a Brainsight neuronavigation system (Rogue Research) to record the hotspot for FDI (Experiment 1) or FCR (Experiment 2) at the beginning of each session to ensure consistent coil placement before and after the perceptual alignment task. Muscle activity was recorded with surface electromyography (EMG) using a belly-tendon montage, with a single common ground electrode over the ulnar styloid process of the target arm. We recorded from FDI and ADM in Experiment 1, and FCR, ECR, and BB in Experiment 2. EMG recordings were amplified (AMT-8; Bortec Biomedical), band-pass filtered (10-1000 Hz), sampled at 5000 Hz, and recorded on a hard drive for subsequent analysis using Signal software (Cambridge Electronic Design Ltd, UK) and MATLAB (Mathworks).

Resting motor threshold (RMT) and the input-output (I/O) curve were measured before and after the perceptual alignment task in each session. We determined RMT for either the FDI (Experiment 1) or FCR (Experiment 2) as the minimum stimulator intensity to evoke MEPs > 50 µV in at least 10 out of 20 trials (Rossini et al. 2015). To assess the I/O curve, monophasic single pulses were delivered at intensities ranging from 90% of RMT up to 200% of RMT, or as high as the subject could tolerate. Intensity order was randomized and inter-pulse interval was 4-6 s. In Experiment 1, we delivered 10 pulses per intensity. In Experiment 2, we delivered 15 pulses per intensity since more proximal MEPs are variable (Carson et al. 2013; Sankarasubramanian et al. 2015). Single trials in which root mean square EMG exceeded 15 microvolts in the 100 ms prior to the TMS pulse were excluded. For each I/O curve (2 per session per muscle per subject), MEP amplitude at each stimulus intensity was calculated and ordered by increasing stimulus intensity. Because not all participants could tolerate TMS intensities over 160% of RMT, and to be consistent with our previous use of this paradigm (Munoz-Rubke et al. 2017), we computed area under the I/O curve over the 90-160% of RMT range for all muscles except BB. This is common practice in both hand and forearm muscles (Meesen et al. 2011; Rossini et al. 2015; Suzuki et al. 2014). However, because BB responses tend to be small when the FCR hotspot is targeted (Carson et al. 2013), for BB we computed area under the I/O curve from 90% of FCR RMT up to the subject’s maximum tolerated intensity. This approach should partially compensate for not targeting the BB hotspot directly. Area under the I/O curve was calculated using the trapezoidal rule. We chose to focus on area under the I/O curve rather than slope, which is also commonly used, because I/O curves may take different shapes for different muscles, especially when slightly different stimulus intensities are captured for each muscle (Carson et al. 2013). In other words, a stimulus intensity of 120% of FCR RMT will yield a different place on the FCR I/O curve than on the BB I/O curve, since the biceps hotspot is somewhat different from the FCR hotspot. Importantly, I/O curves were only compared to other I/O curves obtained from the same muscle.

#### Statistical analysis

To test whether changes in excitability of any of the muscles was related to individual responses to misalignment, we computed a multilevel linear model for percent change (post-alignment task divided by pre) in area under the I/O curve in association with session type (misaligned or veridical). A separate multilevel model was computed for each muscle (FDI and ADM in Experiment 1; FCR, ECR, BB in Experiment 2). Consistent with our previous study (Munoz-Rubke et al. 2017), we computed a reduced model that included only the interaction terms; in other words, the minimum predictors needed to reveal whether individuals’ visual and proprioceptive realignment was related to their change in M1 excitability. The fixed part of each model thus comprised interaction terms of session type with proprioceptive realignment and visual realignment. The models also included a random intercept to account for the repeated-measures design. All models were computed with the lmer4 package, Version 1.1.12 (Bates et al. 2015) of the R programming language (R Core Team 2016).

We checked the normality of variable distributions using the Shapiro-Wilk test. This indicated that area under the I/O curve was not normally distributed for any muscle in any session, and percent change in area under the I/O curve was non-normal for certain muscles in certain sessions: FDI misaligned and ADM veridical session in Experiment 1, and FCR veridical and BB misaligned session in Experiment 2 (all p < 0.05). We therefore log-transformed these variables for all five muscles, in both sessions. This resulted in all variable distributions meeting the assumption of normality. For each muscle, no predictor had a variance inflation factor value greater than 1.5, suggesting the absence of multi-collinearity in the multilevel models. We checked for potential outliers by calculating Cook’s distance (Nieuwenhuis et al. 2012). No data point showed a D value > 1, suggesting no outliers were present for any of the muscles (Cook and Weisberg 1982). To examine the raw I/O curves we computed a repeated measures ANOVA with factors Session (misaligned vs. veridical) and Timepoint (pre- vs. post-alignment task) on the log-transformed area under the I/O curve.

### Experiment 3: Hand and arm proprioception

Each subject completed one session. The session consisted of a modified version of the perceptual alignment task used in Experiments 1 and 2. Subjects were asked to indicate with their right index finger the position of a target related to the left hand or arm.

#### Apparatus and setup

The same apparatus was used as in the first two experiments, with no direct vision of either hand or arm. The indicator finger was again the right index finger, but proprioceptive targets included four points on the left hand and arm: index fingertip, knuckle (anterior first metacarpophalangeal (MP) joint), wrist (anterior midline), and elbow (medial epicondyle). A small square of tape (~5 mm) was placed on each joint to make sure the subject understood which part of each joint to aim for. Participants were instructed to aim for the center of the tape. To avoid fatiguing the target arm, the subject’s left hand and forearm were rested on top of the touchscreen glass, palm down, fingers spread, and slightly to the right of body midline (Fig. 3A).

**Figure 3.**
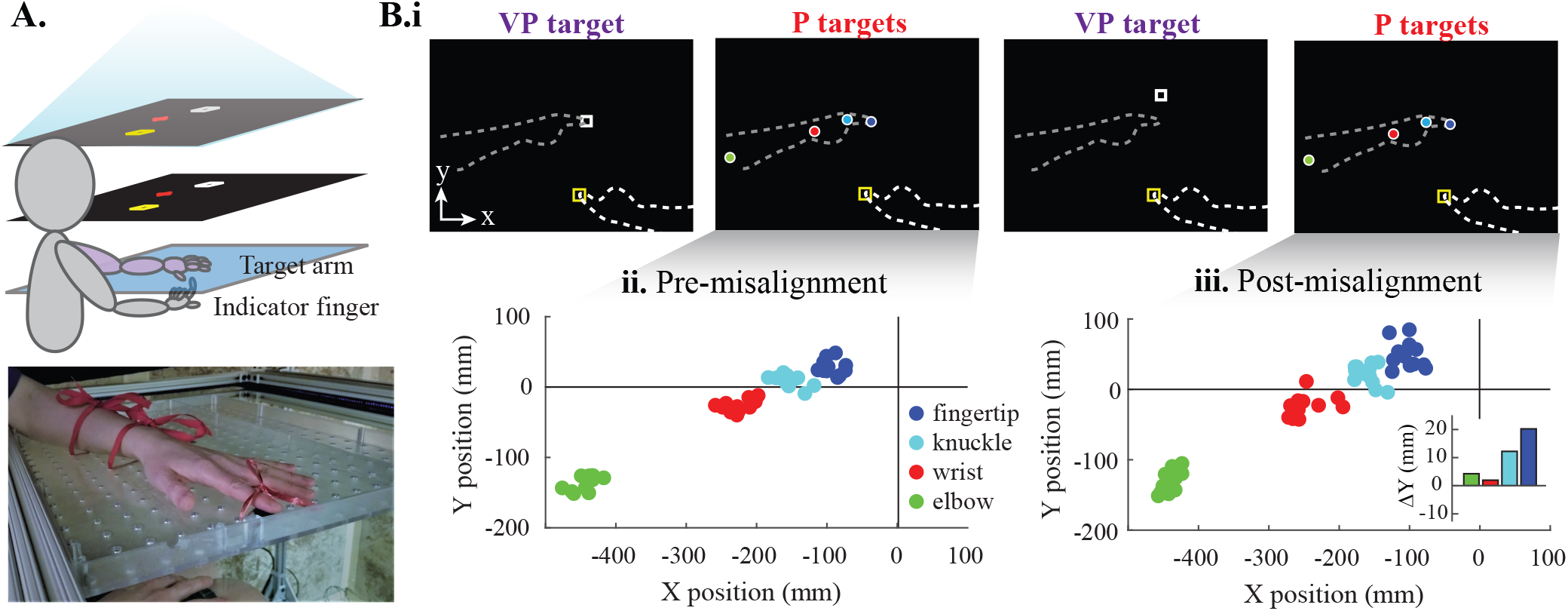
Experiment 3 methods. **A. Apparatus**. Images viewed in the mirror (middle layer) appeared to be in the plane of the touchscreen glass (bottom layer). Subjects pointed with their right (indicator) index finger beneath the glass to where they perceived targets related to their left hand or arm, which rested on top of the glass. **B.i. Session design.** The single session consisted of five blocks of trials: three blocks of proprioceptive targets (P) alternating with two blocks of visuo-proprioceptive targets (VP). For P target blocks, nothing related to the target arm was visible, and subjects were prompted to point to their left index fingertip (blue), knuckle (cyan), wrist (red) or elbow (green). For VP target blocks, only the left index fingertip was a target. Its position was indicated by a white square that was veridical in the first VP block but gradually shifted forward in the second (misaligned) VP block. **ii.** A single subject’s performance on P targets prior to the VP misalignment block. The left index fingertip was always at the origin. **iii.** The same subject’s performance on post-misalignment P targets. Joint estimates shifted forward relative to pre-misalignment (inset), suggesting proprioceptive realignment in response to the VP misalignment block

Prior to the subject placing their left arm in this position, a thin plastic pegboard was secured onto the touchscreen glass. The index fingertip was first placed on a tactile marker on the pegboard, to ensure the fingertip would be at the same location as the visual target. The wrist and elbow were then placed on the pegboard in a way that allowed the fingertip to remain on the tactile marker. Once the subject was comfortable, the left arm was tied to the pegboard at the index finger, the wrist, and the forearm. When positioned, the elbow was bent at about 90 degrees and the wrist was straight. Participants were asked to keep their target arm and hand relaxed and still except during breaks.

The right indicator hand remained below the glass at all times, indicating position estimates on the lower surface of the glass.

#### Procedures

The session consisted of alternating blocks of two types: proprioceptive-only (P targets) and visuo-proprioceptive (VP targets). For each P target block, subjects were asked to indicate their estimate of the left index fingertip, knuckle, wrist, and elbow, 12 times each in random order (48 trials total in the block). No visual information was given about any of the proprioceptive targets during the P block. Each VP block consisted of 42 VP trials in which the subject was asked to indicate their estimate of the left index fingertip with a white box they were told was at the same location.

The session began with a familiarization of each block type, with VP targets displayed veridically during the VP block (Fig. 3Bi). This was followed by a pre-misalignment P block, then a VP block in which misalignment occurred, and finally a post-misalignment P block. During the misalignment VP block, the white box was gradually shifted forward (positive y-dimension) from the left index fingertip, 1.67 mm every other trial, for a maximum misalignment of 35 mm. To maintain circulation and comfort, participants were given a short break between blocks to isometrically contract their left arm, which remained in the same position on the pegboard throughout. They were instructed to do shoulder rolls and to press down with their fingers, hand, and forearm, and then relax. At the conclusion of the experiment, subjects were asked to rate their fatigue, attention, and sleep quality from the previous night on a scale from 1-10.

Each trial began with the right (indicator) hand resting in the subject’s lap. The target arm remained positioned on the pegboard throughout. As in Experiments 1 and 2, subjects were instructed to fixate on the red fixation cross, which appeared at random coordinates within a 10 cm zone around the fingertip target. After an auditory signal, participants indicated their perception by moving their right index fingertip to the lower surface of the glass under which they felt the target was located. As in Experiments 1 and 2, subjects had no direct vision of their arms; received no performance feedback or knowledge of results; had no speed requirements; and were asked to be as accurate as possible. During P target blocks, the specific joint to aim for on each trial was indicated to the subject by text along the top of the reflected display: finger, knuckle, wrist, or elbow.

As with Experiments 1 and 2, we calculated a regression between red cross position and indicator finger endpoints on P fingertip targets. The group average R^2^ was 0.032 ± .019 (mean ± 95% CI), which is consistent with Experiments 1 and 2. We also questioned participants about whether the white box felt like it was on top of their target fingertip. None reported perceiving a forward offset of the white box.

#### Analysis

For each participant, we calculated the change in each joint’s proprioceptive estimate from pre-misalignment to post-misalignment, in the dimension of misalignment (y-dimension; see Fig. 3Bii-iii). To compare realignment at the knuckle, wrist, and elbow to realignment at the fingertip, we computed a correlation between realignment at the fingertip and at each of the other joints. Pearson correlation coefficients were calculated with an α of 0.05, and results are reported after Bonferroni correction.

## Results

### Experiment 1: M1 hand representations

#### Behavioral results

On average, subjects had 13.9 ± 5.2 days between the two sessions (mean ± 95% CI). On a scale of 1-10, subjects rated their quality of sleep the night before at 7.7 ± 0.6 for the misaligned session and 7.6 ± 0.4 for the veridical session. They rated their level of attention at 7.5 ± 0.4 and 7.9 ± 0.4 for the misaligned and veridical sessions, respectively. They rated their fatigue from the experiment at 4.3 ± 0.7 and 3.7 ± 0.8 for the misaligned and veridical sessions, respectively.

Because no misalignment occurred in the veridical session, visual and proprioceptive realignment are expected to be roughly zero on average, across subjects, whereas in the misaligned session we expect them to be above zero on average. In the misaligned session (Fig. 4Ai), participants realigned proprioception 14.5 ± 4.2 mm (mean ± 95% CI), range, and vision 33.9 ± 6.4 mm. Including both visual and proprioceptive realignment, subjects compensated for 48.4 ± 6.1 mm of the 70 mm visuo-proprioceptive mismatch. In the veridical session, subjects realigned proprioception 0 ± 4.9 mm and vision −3.3 ± 5.0 mm. These values are consistent with our previous use of this paradigm (Munoz-Rubke et al. 2017). The mean indicator finger endpoint position was, on average, 23 mm further forward in the misaligned session compared to the veridical session, but the 95% confidence interval of this difference was 31 mm, suggesting any between-session difference in indicator finger movements was highly variable.

**Figure 4.**
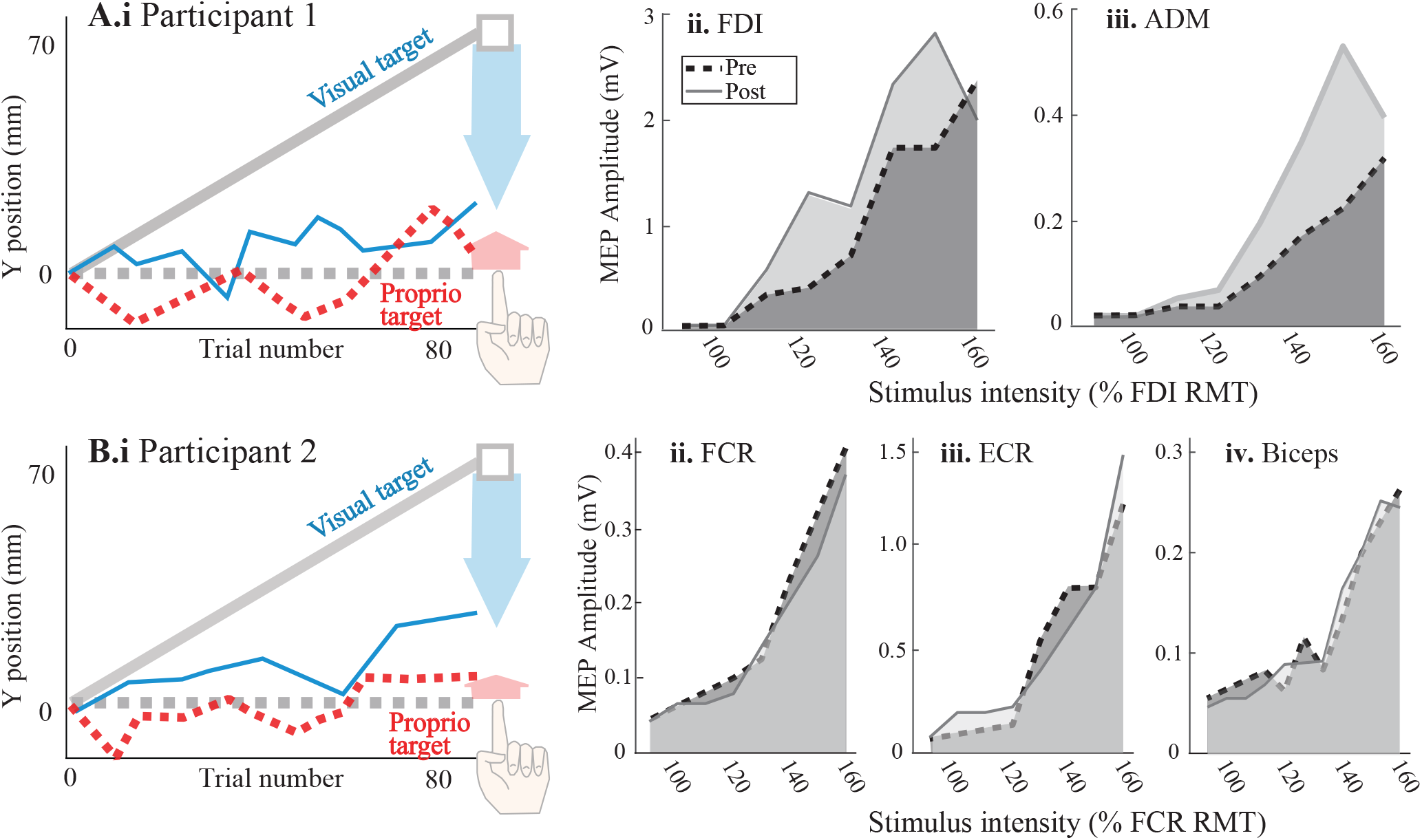
Comparison of two subjects with high visual realignment relative to other subjects. **A.** Experiment 1 participant. **i.** In the misaligned session, this subject realigned proprioception 12.3 mm (red dashed line, red arrow) and vision 57.5 mm (blue line, blue arrow). **ii-iii.** Area under the I/O curve increased after misaligned task 52% for FDI and 90% for ADM. **B.** Experiment 2 participant. **i.** This individual realigned proprioception 10.3 mm (red arrow) and vision 37.7 mm (blue arrow). **ii.** For FCR, area under the I/O curve decreased 11% after the misalignment task. **iii.** ECR area under the I/O curve increased 1%. **iv.** Biceps area under the I/O curve decreased 4%.

#### Neurophysiology

Changes in excitability of the M1 representation of FDI showed realignment modality-specific associations with the misaligned, but not the veridical session. These associations are illustrated with predictor residual plots (Fig. 5Ai), which allows us to show the relationship of one predictor variable with the dependent variable, after statistically controlling for the effect of the other predictor (McElreath 2015). In other words, for the misaligned session only, greater proprioceptive realignment was associated with greater decrease in area under the I/O curve, after controlling for the effect of visual realignment. Conversely, after controlling for the effect of proprioceptive realignment, greater visual realignment was associated with more increase in area under the I/O curve.

**Figure 5.**
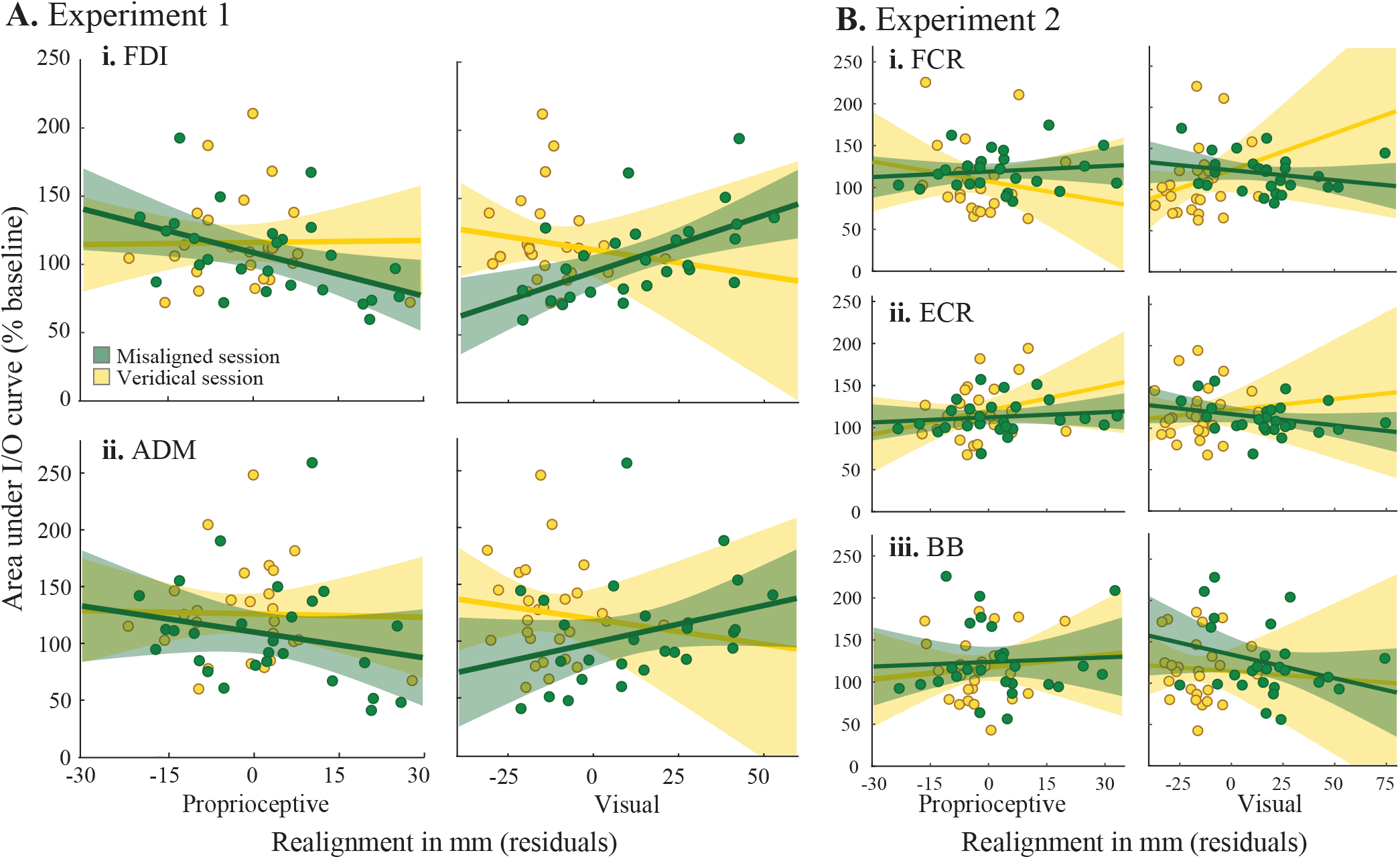
Change in area under the I/O curve (post divided by pre) plotted against predictor residuals, with lines of best fit and corresponding 95% CIs. **A. Experiment 1.** For both FDI (i) and ADM (ii), after statistically controlling for the effect of visual realignment, large proprioceptive realignment in the misaligned session (green) was significantly associated with reduced M1 excitability (left panels). For FDI only, after controlling for the effect of proprioceptive realignment, large visual realignment was associated with increased M1 excitability (right panel). No associations found in veridical session (yellow). **B. Experiment 2.** No significant associations were found for FCR (i), ECR (ii), or BB (iii).

The multilevel model of the association between log percent change in area under the FDI I/O curve and session type (Table 1) indicates a negative association between the misaligned session and proprioceptive realignment (β = −0.01, t_47_ = −2.85, p = 0.004) and a positive association between the misaligned session and visual realignment (β = 0.004, t_47_ = 1.99, p = 0.047). With the dependent variable log-transformed, the betas can be interpreted by exponentiating. The negative association between misaligned session and proprioceptive realignment is equivalent in magnitude to a 1.01% more reduction in FDI area under the I/O curve for every 1 mm more of proprioceptive realignment. The positive association between misaligned session and visual realignment can be interpreted as a 0.37% more increase in FDI excitability for every 1 mm more of visual realignment. No statistically significant associations were observed during the veridical session (p > 0.3). From the perspective of a single subject in the misaligned session, individuals who realigned proprioception substantially generally had a decrease in excitability for the M1 representation of FDI (e.g., Fig. 4Aii). Individuals who realigned vision most extensively generally had an increase in M1 excitability of the FDI representation.

**Table 1.** Experiment 1 multilevel regression results comprising four interaction terms for session type (veridical or misaligned) and realignment type (proprioceptive or visual)

| *Predictors* | FDI |  | ADM |  |
| --- | --- | --- | --- | --- |
| | $\beta$ (CI) | *p* | $\beta$ (CI) | *p* |
| *Fixed parts* |  |  |  |  |
| Intercept | 4.69 (4.60, 4.79) | **<0.001** | 4.74 (4.60, 4.89) | **<0.001** |
| Veridical session : Proprioceptive realignment | 0.0016 (-0.0075, 0.0108) | 0.723 | -0.0005 (-0.014, 0.013) | 0.939 |
| Misaligned session : Proprioceptive realignment | -0.010 (-0.017, -0.003) | **0.004** | -0.013 (-0.023, -0.002) | **0.015** |
| Veridical session : Visual realignment | -0.0040 (-0.013, 0.0049) | 0.375 | -0.004 (-0.017, 0.009) | 0.534 |
| Misaligned session : Visual realignment | 0.0037 (0.0000, 0.0073) | **0.047** | 0.0022 (-0.0031, 0.0075) | 0.418 |
| *Random parts* |  |  |  |  |
| N<sub>ID</sub> | 27 |  | 27 |  |
| Observations | 54 |  | 54 |  |

Columns represent log-transformed percentage of baseline (post divided by pre) in area under the I/O curve. $\beta$ s are presented with their 95% CIs. Boldface identifies statistically significant results.

The multilevel model for ADM also showed a significant association for the misaligned session (Fig. 5Aii). Specifically, a negative association between the misaligned session and proprioceptive realignment was observed (β = −0.01, t_47_ = −2.43, p = 0.015; Table 1). This is equivalent to a 1.27% decrease in ADM area under the I/O curve for every 1 mm more of proprioceptive realignment. The association between the misaligned session and visual realignment was positive, but not statistically significant (β = 0.002, t_47_ = 0.81, p = 0.42). No statistically significant associations were observed during the veridical session (p > 0.5). At the individual participant level, subjects who realigned proprioception substantially in the misaligned session tended to have a greater reduction in M1 excitability of the ADM representation (e.g., Fig. 4Aiii).

Other measures related to M1 neurophysiology and TMS were similar between the misaligned and veridical sessions. For FDI, log-transformed area under the I/O curve showed a main effect of timepoint (F_1,26_ = 5.83, p = 0.023), but no effect of session (F_1,26_ = 1.38, p = 0.25) and no timepoint X session interaction (F_1,26_ = 1.47, p = 0.24). This suggests an increase in excitability for the M1 representation of FDI related to the alignment task, regardless of whether it was the misaligned or veridical task (Fig. 6Ai). For ADM (Fig. 6Aii), log-transformed area under the I/O curve showed no significant effect of timepoint (F_1,26_ = 2.95, p = 0.098), session (F_1,26_ = 0.008, p = 0.92), or interaction (F_1,26_ = 3.13, p = 0.088). On a scale from 1-10, subjects rated the painfulness of TMS at 2.7 ± 0.7 and 2.7 ± 0.6 for the misaligned and veridical sessions, respectively (mean ± 95% CI). FDI RMT was 40.5 ± 2.3% of max stimulator output in the misaligned session, and 41.3 ± 2.3% of max stimulator output in the veridical session.

**Figure 6.**
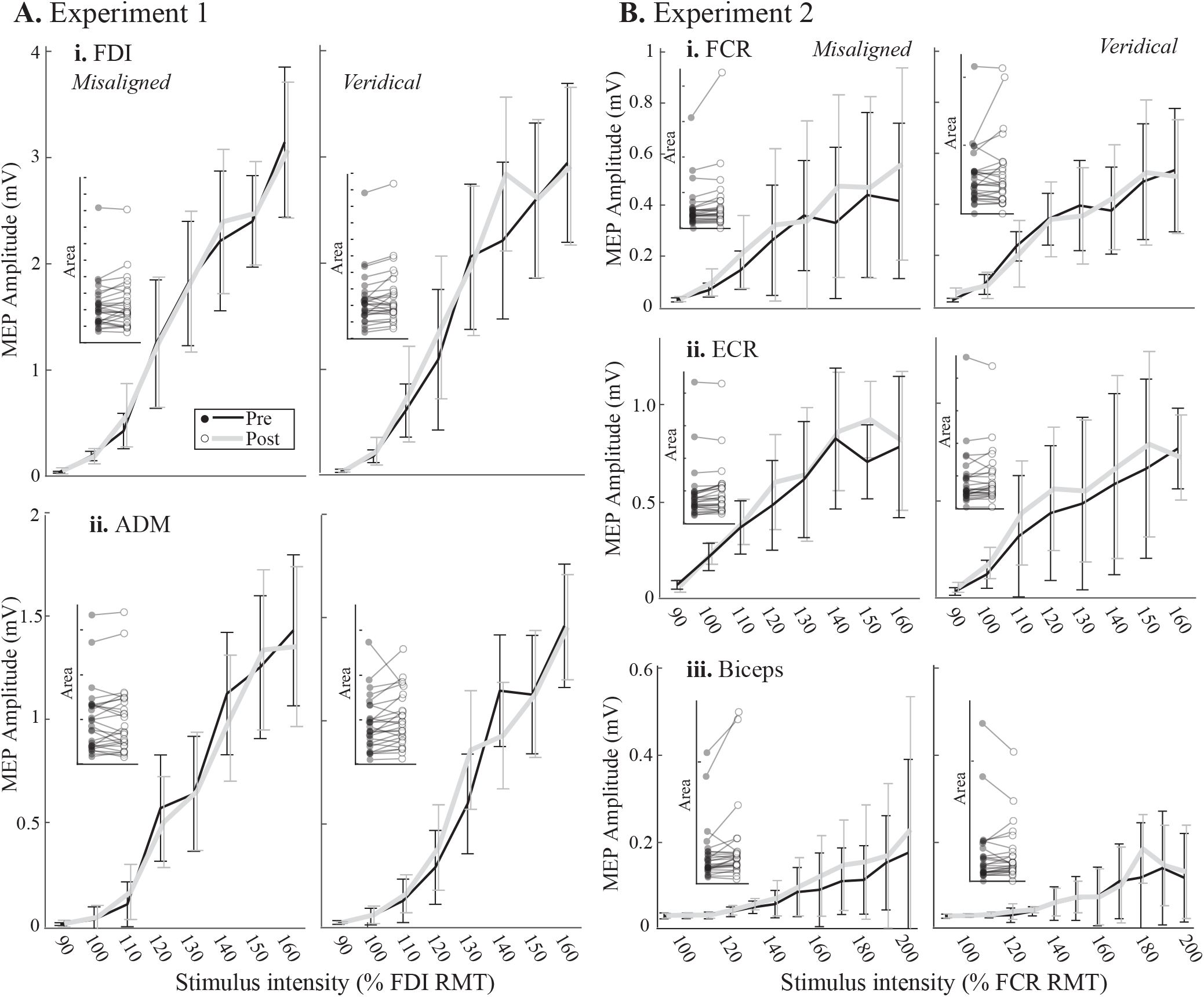
Experiment 1 (A) and 2 (B) input/output curves. Median MEP amplitudes arranged in order of increasing stimulus intensity, pre- and post-alignment task in the misaligned (black lines) and veridical (grey lines) sessions. Error bars represent 95% CI. **Insets:** individual subjects’ area under the input/output curves pre- and post-alignment task. Each notch on the y-axis represents an area of 50.

### Experiment 2: M1 arm representation

#### Behavioral results

Participants had 13.3 ± 5.3 days between the two sessions (mean ± 95% CI). On a scale of 1-10, subjects rated their quality of sleep the night before at 6.9 ± 0.5 for the misaligned session and 7.4 ± 0.6 for the veridical session. They rated their level of attention at 7.2 ± 0.6 and 7.4 ± 0.5 for the misaligned and veridical sessions, respectively. They rated their fatigue from the experiment at 4.3 ± 0.7 and 4.1 ± 0.8 for the misaligned and veridical sessions, respectively.

As in Experiment 1, realignment magnitudes in Experiment 2 were consistent with our previous use of this paradigm (Munoz-Rubke et al. 2017). In the misaligned session, subjects realigned proprioception 9.0 ± 6.3 mm (mean ± 95% CI), and vision 38.2 ± 8.5 mm. Including both visual and proprioceptive realignment, subjects compensated for 47.2 ± 8.5 mm of the 70 mm visuo-proprioceptive mismatch. In the veridical session, subjects realigned proprioception 2.8 ± 3.9 mm and vision 5.1 ± 4.8 mm. The mean indicator finger endpoint position was, on average, 21 mm further forward in the veridical session compared to the misaligned session, but the 95% confidence interval of this difference was 62 mm, suggesting any between-session difference in indicator finger movements was highly variable.

#### Neurophysiology

Results do not indicate any realignment modality-specific associations with the misaligned session for FCR (Fig. 5Bi), ECR (Fig. 5Bii), or BB (Fig. 5Biii). In other words, neither visual nor proprioceptive realignment was significantly associated with log percent change in area under the I/O curve for any of the three muscles recorded in Experiment 2; this is seen even in subjects who realigned the most (Fig. 4B). Full multilevel model results are presented in Table 2.

**Table 2.**
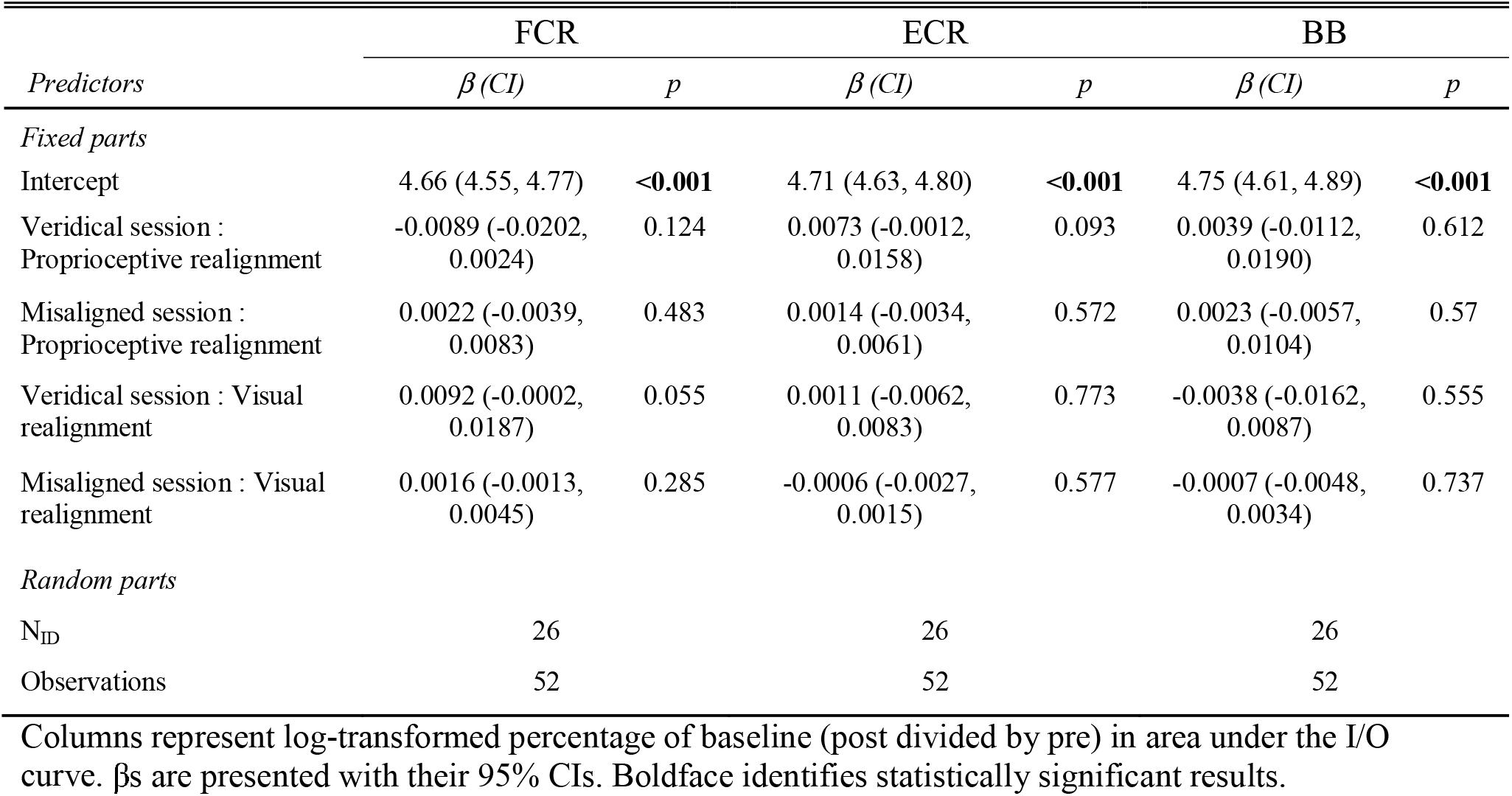
Experiment 2 multilevel regression model results comprising four interaction terms for session type (veridical or misaligned) and realignment type (proprioceptive or visual)

| *Predictors* | FCR |  | ECR |  | BB |  |
| --- | --- | --- | --- | --- | --- | --- |
| | $\beta$ (CI) | *p* | $\beta$ (CI) | *p* | $\beta$ (CI) | *p* |
| *Fixed parts* |  |  |  |  |  |  |
| Intercept | 4.66 (4.55, 4.77) | **<0.001** | 4.71 (4.63, 4.80) | **<0.001** | 4.75 (4.61, 4.89) | **<0.001** |
| Veridical session :<br>Proprioceptive realignment | -0.0089 (-0.0202,<br>0.0024) | 0.124 | 0.0073 (-0.0012,<br>0.0158) | 0.093 | 0.0039 (-0.0112,<br>0.0190) | 0.612 |
| Misaligned session :<br>Proprioceptive realignment | 0.0022 (-0.0039,<br>0.0083) | 0.483 | 0.0014 (-0.0034,<br>0.0061) | 0.572 | 0.0023 (-0.0057,<br>0.0104) | 0.57 |
| Veridical session : Visual<br>realignment | 0.0092 (-0.0002,<br>0.0187) | 0.055 | 0.0011 (-0.0062,<br>0.0083) | 0.773 | -0.0038 (-0.0162,<br>0.0087) | 0.555 |
| Misaligned session : Visual<br>realignment | 0.0016 (-0.0013,<br>0.0045) | 0.285 | -0.0006 (-0.0027,<br>0.0015) | 0.577 | -0.0007 (-0.0048,<br>0.0034) | 0.737 |
| *Random parts* |  |  |  |  |  |  |
| N<sub>ID</sub> | 26 |  | 26 |  | 26 |  |
| Observations | 52 |  | 52 |  | 52 |  |

Columns represent log-transformed percentage of baseline (post divided by pre) in area under the I/O curve. $\beta$ s are presented with their 95% CIs. Boldface identifies statistically significant results.

In contrast, all three muscles showed a significant increase in log area under the I/O curve from pre- to post-alignment task (main effect of timepoint: F_1,25_ = 5.82, p = 0.023 for FCR; F_1,25_ = 10.7, p = 0.003 for ECR; and F_1,25_ = 9.30, p = 0.005 for BB). This suggests the M1 representation of all three muscles increased in excitability after the task, regardless of whether it was the misaligned or veridical session (Fig. 6B). FCR had no effect of session (F_1,25_ = 0.02, p = 0.88) and a timepoint X session interaction that did not rise to the level of significance (F_1,25_ = 3.53, p = 0.072). ECR had no session effect (F_1,25_ = 1.97, p = 0.17) and no interaction (F_1,25_ = 0.33, p = 0.57). BB had no significant effect of session (F_1,25_ = 4.05, p = 0.055) and no timepoint X session interaction (F_1,25_ = 0.40, p = 0.53). This suggests that biceps excitability in the two sessions was slightly different, both before and after the alignment task, but did not change differently from pre- to post-alignment task in one session compared to the other.

On a scale from 1-10, subjects rated the painfulness of TMS at 1.8 ± 0.3 and 2.0 ± 0.6 for the misaligned and veridical sessions, respectively (mean ± 95% CI). FCR RMT was 41.4 ± 2.4% of max stimulator output in the misaligned session, and 40.8 ± 2.1% of max stimulator output in the veridical session.

### Experiment 3: Hand and arm proprioception

After a 35 mm visuo-proprioceptive misalignment at the left index fingertip, some subjects showed a marked forward shift in in their proprioceptive estimate of that finger (e.g., Fig. 3Bii-iii). However, on average proprioceptive realignment was 1.9 ± 4.4 mm (mean ± 95% CI) at the fingertip. Fingertip realignment was closely matched by proprioceptive realignment at the knuckle (Fig. 7A), with a correlation of 0.78 (p = 0.001). Proprioceptive realignment at the wrist was not related to realignment at the fingertip (r = 0.25, p = 0.38; Fig. 7B), and neither was proprioceptive realignment at the elbow (r = −0.37, p = 0.19; Fig. 7C). On a scale of 1-10, the 14 subjects rated their quality of sleep at 6.9 ± 0.7, their level of attention at 6.6 ± 1.1, and their level of fatigue at 4.4 ± 1.2.

**Figure 7.**
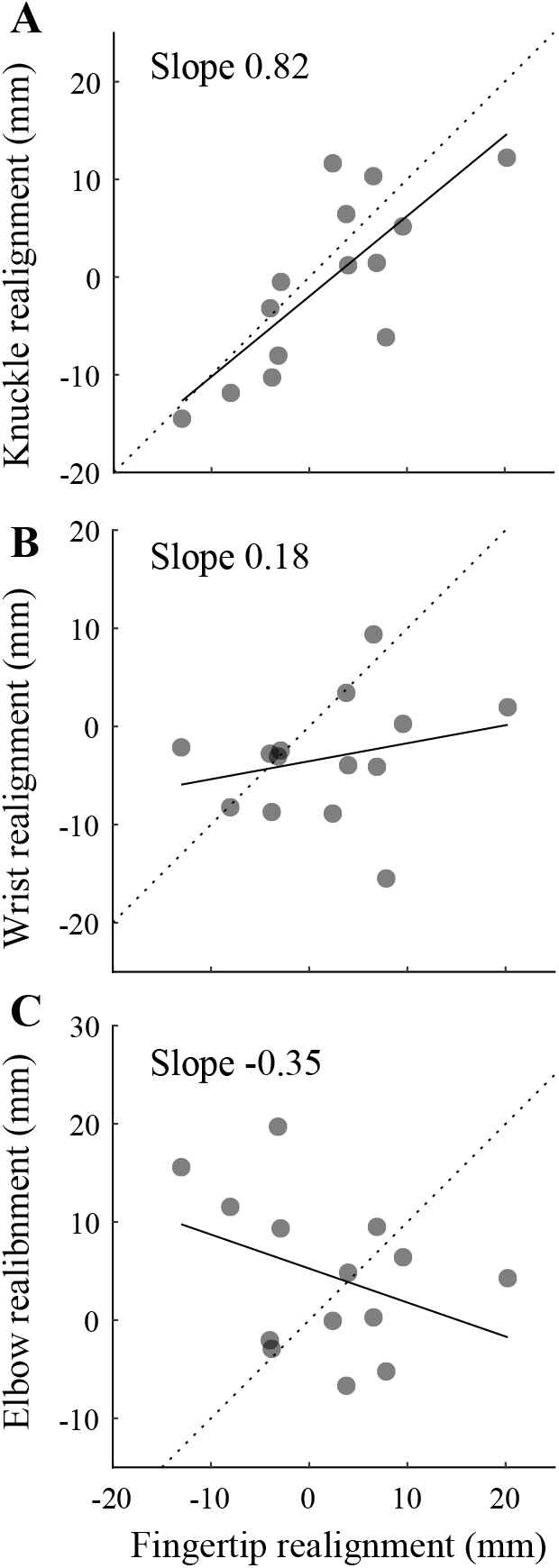
Experiment 3 results. Realignment in proprioceptive estimates of the knuckle (**A**), wrist (**B**), and elbow (**C**) targets, compared to proprioceptive realignment at the left index fingertip. Dotted line represents a 1:1 relationship between realignment at the two joints. Each circle represents one subject.

## Discussion

Here we asked whether visuo-proprioceptive realignment at the fingertip affects only the brain’s representation of the misaligned finger (somatotopically focal), or whether changes extend to other parts of the hand and arm that would be needed to move the misaligned finger (somatotopically broad). Results from three experiments support the first option (Fig. 1B): at both the level of M1 neurophysiology and conscious perception, hand regions near the misaligned fingertip were affected, but further parts of the limb were not.

### M1 representation of the misaligned index finger showed realignment-specific changes

Results from FDI in Experiment 1 replicate our previous findings (Munoz-Rubke et al. 2017): Area under the I/O curve decreased in association with proprioceptive realignment, and increased in association with visual realignment in the misaligned session. In other words, M1 excitability changes were specific to the sensory modality of realignment. This is consistent with a neurological model in which changes in visuo-proprioceptive integration are reflected in M1 via altered connections from brain areas traditionally considered unisensory. While multisensory influences on M1 have rarely been studied, proprioceptive training (Wong et al. 2012) and stimulation (Carel et al. 2000; Lewis and Byblow 2004) are known to affect movement and motor system plasticity, respectively. Our findings are also consistent with the rubber hand illusion (RHI). The RHI involves a discrepancy between the seen fake arm and the felt real arm that creates the illusion of body ownership over the fake arm. This paradigm is thought to involve proprioceptive realignment (Butler et al. 2017) and has been linked to reduced M1 excitability (Della Gatta et al. 2016; Isayama et al. 2019). Similarly, in our results, proprioceptive realignment was associated with reduced M1 excitability for FDI.

The modality-specific associations for FDI representation were present in the misaligned, but not the veridical (control) session, suggesting a specific connection to visuo-proprioceptive learning. The target hand performed the same actions throughout both sessions—touching a stationary tactile marker or resting in the subject’s lap—ruling out movement-related explanations for the misalignment effects. Subjects received no movement error feedback or knowledge of results in either session, to prevent trial-and-error motor adaptation.

### M1 representation of other muscles did not show the same pattern

Modality-specific associations with M1 excitability were somatotopically focal. Like FDI, ADM excitability was negatively associated with proprioceptive realignment in the misaligned session. The ADM association with visual realignment, while positive like FDI, was not significant. In a demographically similar group of healthy young adults, FCR, ECR, and BB showed no significant associations in the misaligned session. In other words, the pattern of associations observed for the misaligned index finger was partially present for the little finger, but absent for the arm muscles, apparently diminishing with distance from the misaligned finger. This suggests that visuo-proprioceptive realignment has localized neurophysiological effects, not broadly generalized to motor representations of body parts involved in moving the misaligned finger. While the present study is the first, to our knowledge, to test M1 representations in the context of multisensory perceptual learning, our somatotopically focal results are consistent with the effects of peripheral somatosensory stimulation on M1. For example, vibration of one muscle increases M1 excitability for that muscle, but decreases it for neighboring muscles (Rosenkranz and Rothwell 2004).

TMS measures in proximal muscles are more variable than in distal muscles (Brasil-Neto et al. 1992; Harris-Love et al. 2007; Sankarasubramanian et al. 2015). However, others have found significant changes in proximal I/O curve areas after interventions such as paired associative stimulation (Carson et al. 2013). Indeed, Carson et al. (2013) used 6 TMS pulses per intensity while we used 15, to better control for variability in proximal responses and for variability linked to non-optimal targeting (i.e., targeting FCR for the BB I/O curve) (Brasil-Neto et al. 1992). In any case, all three arm muscles had an increase in excitability after the alignment task, regardless of session, suggesting the measure was sensitive enough to detect change. Biceps responses were consistent with others who targeted the FCR hotspot (Carson et al. 2013), but small amplitudes pre-alignment task raise the possibility of a floor effect. In other words, perhaps subjects who might otherwise have shown a decrease in BB excitability (i.e., realigned proprioception a lot, if the FDI pattern is followed) started out with such low BB responses that no reduction could be detected. However, of the two subjects whose pre-misalignment biceps responses were on the low end of results in (Carson et al. 2013), neither had a high proprioceptive realignment magnitude. Therefore, a floor effect is unlikely to be a factor in the biceps results.

### Perceptual changes are also somatotopically focal

In Experiment 3, we found that changes in conscious perception of index fingertip position were closely related to proprioceptive estimates of the knuckle (first MP joint), but not the wrist or elbow. In other words, distortion in the body’s representation is localized: If the index finger and knuckle feel like they are further away, perception of wrist and elbow positions do not change to match. The RHI can also distort the body representation, for example with an extra-long arm (Kilteni et al. 2012) or mismatched arm segment lengths (Perez-Marcos et al. 2018). Others have suggested that the brain strives to minimize distortion in the body image, based on evidence that the RHI affects perceived elbow angle as well as hand position (Butz et al. 2014). However, the fake “hand” normally comprises both a hand and a forearm (Butz et al. 2014), so there is a visuo-proprioceptive mismatch not only between the real and fake hand, but also the real and fake forearm, which should be sufficient to drive proprioceptive realignment at the elbow. A version of the RHI more analogous to the present study might require covering all parts of the fake hand and forearm except the index finger, then testing perceived positions of multiple hand and arm joints.

Although the Experiment 3 perceptual results are consistent with the somatotopically focal results in M1, some differences in the alignment task merit consideration. As we have done previously, the target finger in Experiments 1 and 2 was placed on one of two tactile markers beneath the glass for P and VP trials, and otherwise rested in the lap. Visuo-proprioceptive misalignment was 70 mm, and proprioceptive realignment at the fingertip was around 10 mm. Misalignment in Experiment 3 was only 35 mm, so we might have expected proprioceptive realignment at the fingertip to be around 5 mm. Instead, it averaged only 2 mm. This is likely due to static positioning of the target arm, which rested fingertips-to-elbow on top of the glass throughout the task and was tied in place. Thus, subjects had substantial tactile feedback telling them nothing was moving forward, contradicting the visual information. An additional limitation of this approach is that visual realignment cannot be assessed if the target arm (proprioception) is not removed from the workspace. Further study of this paradigm, perhaps with other target arm configurations and measurement of visual estimates, could improve our understanding of the changes in body perception associated with visuo-proprioceptive realignment.

### Implications for sensorimotor function

Our findings of somatotopically-focal effects of visuo-proprioceptive realignment on both motor neurophysiology and body perception are consistent with a tight relationship between sensory and motor systems. The effects of motor learning on sensory neurophysiology (Mirdamadi and Block 2020; Nasir et al. 2013; Vahdat et al. 2011) and perception (Henriques and Cressman 2012; Ostry et al. 2010; Salomonczyk et al. 2012; Wong et al. 2012) have been studied in some detail, at least with regard to proprioception. However, the specific effects of perceptual learning on the motor system have rarely been examined directly. Existing examples include work showing that somatosensory training affects functional connectivity with frontal motor regions (Vahdat et al. 2014); in addition, perceptual changes linked to the RHI have been found to affect certain voluntary movements (Botvinick and Cohen 1998; Kammers et al. 2009). The present study builds on this literature by examining the somatotopic effects of a multisensory perturbation on both M1 neurophysiology and body perception.

Our findings raise the possibility that perceptual learning may not generalize to other body parts as motor learning does. Rather, visuo-proprioceptive misalignment appears to create a localized distortion in the multisensory body representation, with correspondingly local changes in the motor cortex representation. In contrast, trial-and-error motor adaptation transfers preferentially from proximal to distal effectors (Krakauer et al. 2006), and motor skill learning may transfer symmetrically within a limb (Rajan et al. 2019). Interestingly, this is consistent with evidence that within-limb representations overlap in M1 more than in S1 (Cunningham et al. 2013) and supports the targeting of proximal muscles for motor rehabilitation (Carson et al. 2013; Fujiyama et al. 2014). Further research will be needed to understand why perceptual learning does not generalize like motor learning, and whether multisensory rehabilitation techniques such as VR (Ekman et al. 2018; Keshner and Fung 2017) and mirror training (Bolognini et al. 2015) should have a somatotopically focal component.

## Acknowledgments

This material is based upon work supported by the National Science Foundation under Grant No. 1753915 to HJB. The authors would like to acknowledge Dr. Felipe Munoz-Rubke and Ms. Stephanie Dickinson for their advice on the statistical analysis of Experiments 1 and 2.

## Notes

**Conflict of Interest statement:** The authors declare no competing financial interests.

### Competing Interest Statement

The authors have declared no competing interest.

